# Unipept in 2026: A Unified Web Platform for Comparative Metaproteomics Analysis

**DOI:** 10.64898/2026.11.23.768510

**Authors:** Tibo Vande Moortele, Simon Van de Vyver, Ben-Björn Binke, Tim Van Den Bossche, Lennart Martens, Pieter Verschaffelt, Peter Dawyndt, Bart Mesuere

## Abstract

Metaproteomics studies the collective proteome of microbial communities to characterize their taxonomic composition and functional activity. Unipept has long supported such analyses, but organizing samples into projects, building targeted reference databases, and comparing samples across a study were only possible in a standalone desktop application. Here we present a major update that moves these workflows into the web application, so a complete multi-sample study can be assembled, analyzed, and shared without local installation. Samples can be compared both taxonomically and functionally, the latter by mapping enzymes onto metabolic pathways with the integrated PathwayPilot. The analyses also became more informative. The Peptonizer2000, now accelerated by the new NORI inference engine, adds confidence scores to taxonomic and functional results, and an optional filter removes common contaminants before analysis. The Unipept API itself was also reworked, with an optimized index and flexible memory loading that roughly doubled query throughput and lowered the memory needed for deployment. We demonstrate these capabilities on six datasets of a defined microbial community.

## Introduction

Metaproteomics, the large-scale study of the collective proteome of microbial communities, has become a standard approach for investigating the taxonomic composition and functional activity of complex ecosystems^1–4^. Unipept^5–8^ (unipept.ugent.be), first introduced in 2012, supports such analyses by matching identified peptides against the UniProtKB^9^ database and aggregating their taxonomic annotations using a lowest common ancestor (LCA) algorithm^5^, together with functional annotations, giving fast, interactive insight into microbial community composition. Over more than a decade of development, Unipept has expanded into an ecosystem of tools built on a common API: a web application, a desktop application, and a command-line interface.

In our previous manuscript, Unipept in 2024^10^, we described how Unipept’s long-standing relational database was replaced by a Sparse Suffix Array (SSA) index, and how its API server was rewritten from Ruby-on-Rails to the performance-oriented Rust language. That update also lifted the restriction to tryptic peptides, adding support for semi-tryptic, non-tryptic, and missed-cleavage peptides, and extended Unipept to fields such as immunopeptidomics. Because these changes were made at the API level, the entire ecosystem benefited.

Despite these improvements, the most advanced workflows were only possible in the standalone Unipept Desktop application^6^: organizing multiple samples into projects, constructing targeted protein reference databases, and comparing samples across a study. The Desktop application builds its own protein reference databases independently of the hosted index. This process can be time-consuming and can require substantial local resources. When it was first developed, browser technology could not support such complex client-side workflows, but modern browsers have since matured. Indeed, file-system APIs to access the local device’s storage, IndexedDB to store more data on the client, and performant Web Workers for background computation have now made it feasible to deliver rich, desktop-class applications entirely within the browser.

Here, we provide an overview of developments across the Unipept ecosystem since the 2024 update. The web application now supports project-based organization of multi-sample studies, selection of targeted protein reference subsets, and comparative taxonomic analysis through interactive heatmaps and barplots. Project export allows users to share their data, settings, and results for inspection in another browser session.

The update also integrates probabilistic inference and pathway visualization into these workflows. The Peptonizer2000^11^, accelerated by the NORI inference engine^12^, provides confidence scores for taxonomic and functional results through client-side inference. The integration of PathwayPilot^13^ enables users to explore and compare functional annotations in the context of KEGG metabolic pathways^14^. Alongside these additions, optimizations to the API improve query throughput and reduce deployment memory requirements, while a rewritten command-line interface supports automated workflows. **Table 1** summarizes the input, operations, and output of each of these tools. We demonstrate the updated web functionality using six SIHUMIx datasets from the Critical Assessment of MetaProteome Investigation (CAMPI) study^15^ and evaluate the performance of the revised backend.

**Table 1:** Input, operations, and output of the Unipept tools described in this work. The rows follow the order of the Methods.

| **Tool** | **Input** | **Operations** | **Output** |
|---|---|---|---|
| Project management | Samples and analysis settings | Groups, stores and exports samples | .unipept project file |
| Custom database construction | Taxa, protein accessions or reference proteomes | Restricts protein matches to the selection | Targeted reference for a project |
| Taxonomic comparison | Samples in a project | Compares species-level assignments | Heatmap and barplot |
| Functional comparison (PathwayPilot) | EC numbers of the samples | Maps them onto KEGG pathways | Pathway maps and data |
| Peptonizer2000 with NORI | Peptides of a sample | Probabilistic inference | Taxa and functions with confidence scores |
| Unipept API | Peptide sequences | SSA index lookup | Taxonomic and functional annotations |
| Unipept CLI | Peptide files or standard input | Queries the Unipept API | Annotations in the existing output formats |

## Methods

### Web-based project and custom database management

Project management and custom databases were, until now, available only in the Unipept Desktop application. Bringing both into the web application unifies Unipept’s functionality and removes the need for a local installation.

#### Project management

A project groups one or more samples, together with their analysis settings and results, and is created and managed directly in the browser. All project data is stored client-side using the browser’s IndexedDB API. Changes are saved automatically, so work is preserved even if the browser tab is closed unexpectedly, and a session can be resumed later.

To support collaboration and archiving, projects can now be exported. Project data, settings, databases and results are packaged into a single self-contained *.unipept* file that can be shared and re-imported into Unipept’s web application on any machine. These files are specific to the web application and are not interchangeable with the folder-based project structure used by the Desktop application, as the two tools manage state differently.

#### Custom database construction

By default, Unipept searches the entire UniProtKB-derived index. A custom database restricts this search to a chosen subset of proteins, defined in the web interface by selecting NCBI^16^ taxon identifiers, UniProtKB protein accessions, or reference proteome identifiers. Selecting a taxon also includes every taxon below it, so selecting a species, for example, also covers its strains. Analyses can optionally apply a contaminant filter, which excludes the proteins on the cRAP list (https://www.thegpm.org/crap/index.html) from the matches, so that frequent contaminants such as keratins and trypsin do not affect the results.

Once a custom database has been defined, Unipept’s API server translates the selected taxa, protein accessions, and reference proteomes into a single list of UniProtKB proteins. This list then filters the regular search. For each peptide, Unipept first retrieves every UniProtKB protein that contains it, as in a standard analysis. Proteins outside the custom database are then discarded, and the lowest common ancestor and functional annotations of the peptide are derived from the proteins that remain. A custom database therefore does not require a new search index to be built. This contrasts with the Desktop application, where building a custom reference database is a self-contained process that can take hours to days and requires substantial local disk space and memory.

### Comparative analysis framework

Comparative analysis reveals the differences between samples that single-sample analysis cannot show. Grouping samples into projects makes this possible, both taxonomically and functionally.

#### Taxonomic comparison

A redesigned interactive heatmap shows the relative abundance of each taxon across the selected samples, so individual taxa can be inspected and followed across samples. A comparative barplot complements it with a higher-level view of how community composition shifts between samples, and which taxa dominate.

#### Functional comparison

PathwayPilot provides a pathway-centric view of a sample’s functional activity, mapping functional annotations from matched peptides onto KEGG metabolic pathways. Because it is integrated, a functional comparison can be launched without leaving the web application or re-uploading data. On these interactive pathways, functional profiles can be compared directly, rather than through lists of Enzyme Commission (EC) numbers^17^, Gene Ontology (GO) terms^18,19^, or InterPro entries^20^, placing functional differences in their metabolic context.

### Probabilistic inference with the Peptonizer

A standard Unipept analysis reports how many peptides support each taxon and function. The Peptonizer2000 instead weighs the evidence from all peptides in a sample to assign every result a confidence score. A new inference engine makes it fast enough to run in the browser.

#### Confident taxonomic inference

Whereas the standard LCA approach assigns each peptide independently, the Peptonizer combines the peptide evidence into a single probabilistic model and reports a ranked list of taxa, each with a confidence score. It reuses Unipept’s existing infrastructure, retrieving the required evidence through the same API that drives the regular analysis. It respects the active custom database, so inference can be scoped to a targeted subset of proteins.

#### A faster inference engine

The original Peptonizer2000 ran as an offline pipeline outside the browser. NORI reimplements the same belief-propagation algorithm^21,22^ as a Rust library that is compiled to WebAssembly and runs in Web Workers, so the inference runs client-side, alongside the rest of the analysis.

#### Confidence scores for functional analysis

The same engine also makes confidence scoring practical for functional annotations. Alongside the peptide counts Unipept reports for a variety of ontologies, each functional annotation now carries a confidence score from the same inference model. The score lets functional results be ranked and filtered by how well the peptides support them.

### A faster, more reliable backend

Every capability just described runs against the same Unipept API, computing results on demand rather than reading them from precomputed tables. The computational load therefore falls on the API. Comparative analysis, which loads many samples at once, is the most demanding case. The improved API addresses this in two ways, through an optimized index and flexible index loading.

First, the suffix-array index and its lookup path were optimized with a k-mer seed table, cross-query prefetching, and reduced per-match work. The higher throughput keeps interactive comparison of many samples responsive.

Second, the index can be loaded in three ways that differ in how much of it is held in memory. Preloaded mode keeps the whole index in memory, while memory-mapped (mmap) mode leaves residency to the operating system, so it needs no large fixed allocation. Because mmap lookups can block on page faults, the server runs more worker threads than CPU cores, keeping the processor busy while other threads wait for pages to load. In production, Unipept uses a hybrid approach, preloading every structure except the suffix array. Each instance then requires far less memory than a full preload. The lower footprint lets each instance run on more modest hardware and keeps the service stable when memory is tight or a process restarts.

### A more reliable and reproducible CLI

The Unipept command-line interface (CLI) provides programmatic access to the Unipept API for use in other data-processing pipelines. It has been completely rewritten in TypeScript, replacing the previous Ruby implementation and aligning the CLI with the rest of the Unipept codebase. Its static type system catches many error classes before a command runs. The rewrite preserves full functional parity with the Ruby version, supporting all commands and output formats, so existing pipelines continue to work unchanged.

The CLI can now be run without prior installation through npx, and pinning the version lets a pipeline lock onto an exact release. The CLI reports the UniProtKB release used by the API, allowing users to record the reference-data version alongside the CLI version. A consistent set of global options now handles input and output across all commands. Because every command runs against the same API, the CLI also inherits the backend improvements described above. Together with broader automated test coverage, these changes make it more reliable and its results easier to reproduce.

### AI-assisted development

The software was designed and developed by the authors. Claude Code (Anthropic) was used as a coding assistant to implement parts of the code under the direction of the developers. All code, including AI-assisted contributions, was reviewed multiple times by a developer before inclusion, and the authors remain responsible for its correctness.

## Results

### The web-based project workflow

To demonstrate the new web-based workflow, we analyzed the SIHUMIx^23^ dataset from the CAMPI study^15^ (PRIDE^24^ accession PXD023217), part of the Metaproteomics Initiative^25,26^. SIHUMIx is a simplified human intestinal microbiota model comprising eight bacterial species: *Anaerostipes caccae, Bacteroides thetaiotaomicron, Bifidobacterium longum, Blautia producta, Clostridium butyricum, Clostridium ramosum, Escherichia coli, and Lactobacillus plantarum*.

To cover a range of laboratory workflows, we used the six SIHUMIx datasets that the CAMPI study selected as representative of its fractionation methods and MS instruments (S03, S05, S07, S08, S11, and S14). For each dataset, the X!Tandem^27^ search results against the UniProtKB reference proteome database (SIHUMIx_REF) were retrieved from PRIDE, pooling the four fractions of S11 and the three replicates of S14. Peptide-spectrum matches were filtered at a target-decoy false discovery rate (FDR) below 1% (0.32 – 0.58% per sample), using the threshold applied in CAMPI. Decoy matches were removed, as were spectra whose best-scoring hits were tied between different peptides, yielding between 13 391 (S07) and 35 349 (S11) unique peptides per sample (**Table S1**).

Because the taxonomic composition of SIHUMIx is known, we selected its eight constituent species in Unipept’s taxonomy browser to build a targeted protein reference database. Two species are listed under reclassified names in UniProtKB, *Thomasclavelia ramosa* (formerly *Clostridium ramosum*) and *Lactiplantibacillus plantarum* (formerly *Lactobacillus plantarum*). These taxa were resolved into a set of roughly 105 000 UniProtKB protein identifiers, which then restricted the protein matches in every downstream analysis.

We created a new project in the web application and added the six samples. All samples were analyzed against UniProtKB release 2026_02 with the “Equate I and L” and “Filter duplicate peptides” options enabled and the contaminant filter disabled. Missed cleavages are always taken into account by Unipept. Because the peptides were identified against UniProt reference proteomes, all but three of them matched UniProtKB, and between 99.22% and 99.55% per sample were matched by the eight-species database (**Table S1**). Restricting the reference to the known community therefore discarded very few matches while concentrating the analysis on its members. The complete project is provided as a single .unipept file (**File S1**) that can be loaded into Unipept to inspect the analyses.

### Confident taxonomic and functional analysis

To test whether the Peptonizer2000 recovers the community without prior knowledge, we ran it against the full UniProtKB instead of the eight-species database. No search-engine scores were provided, so default peptide scores were used. In all six samples, *E. coli*, *B. thetaiotaomicron*, *C. butyricum*, *T. ramosa*, *A. caccae*, and *B. longum* reached a confidence of 1.00. *L. plantarum*, the least abundant member, scored at least 0.97 in four samples. *B. producta* never exceeded 0.48, whereas the closely related *Blautia pseudococcoides* scored at least 0.94 in every sample. In S03, the remaining taxa among the 20 highest-scoring species scored 0.86 or lower, and all but *Longibaculum muris* (0.15) belong to genera of community members. *Homo sapiens* reached 1.00 in three samples and *Sus scrofa* in one, most likely reflecting laboratory contaminants such as keratins and porcine trypsin. Enabling the contaminant filter removed *H. sapiens* from the 20 highest-scoring species in all samples and lowered *S. scrofa* to 0.61.

The Peptonizer2000 also scores functional annotations. For each GO term, EC number, and InterPro entry, it reports an estimated probability next to the peptide count. Taking S03 as an example, 274 of the 1 827 EC numbers had a probability above 0.9, which narrows the functional profile to the reactions best supported by the data. The taxonomic inference for S03 took about 10 s in the browser, including the setup of the Peptonizer, whereas the original offline pipeline needed about 3.5 minutes. Probabilistic analysis can therefore be run within the same interactive session as the rest of the workflow.

### Comparative analysis of a defined community

An interactive heatmap and a comparative barplot show the species-level profiles of all six samples side by side (**Figure 1**). *B. thetaiotaomicron* contributed the largest proportion in every sample (52.4 – 69.5%), while *B. producta* (12.4 – 15.9%) and *E. coli* (8.8 – 11.9%) were present at similar levels across samples. All eight species were detected in every sample, although *C. butyricum*, *B. longum*, and *L. plantarum* each accounted for less than 0.4%. These profiles agree with CAMPI, which also found *B. thetaiotaomicron* to be the most abundant species and *E. coli* at 10 – 13%, albeit based on spectral counts. In contrast, CAMPI barely detected *B. producta*, even though about 15% of its peptides were assigned to the genus *Blautia*. At the time, no UniProtKB reference proteome was available for *B. producta*, so the CAMPI reference database contained several *Blautia sp.* proteomes instead, including that of strain YL58, now the type strain of *B. pseudococcoides*^28^. We observed the same pattern against the full UniProtKB, where 11 – 14% of the peptides were assigned to *Blautia* or one of its species, but few of these were resolved to species level, consistent with the Peptonizer results above. The eight-species database assigns these peptides to *B. producta*, the only *Blautia* species it contains, a gain in resolution that relies on prior knowledge of the community.

**Figure 1:**
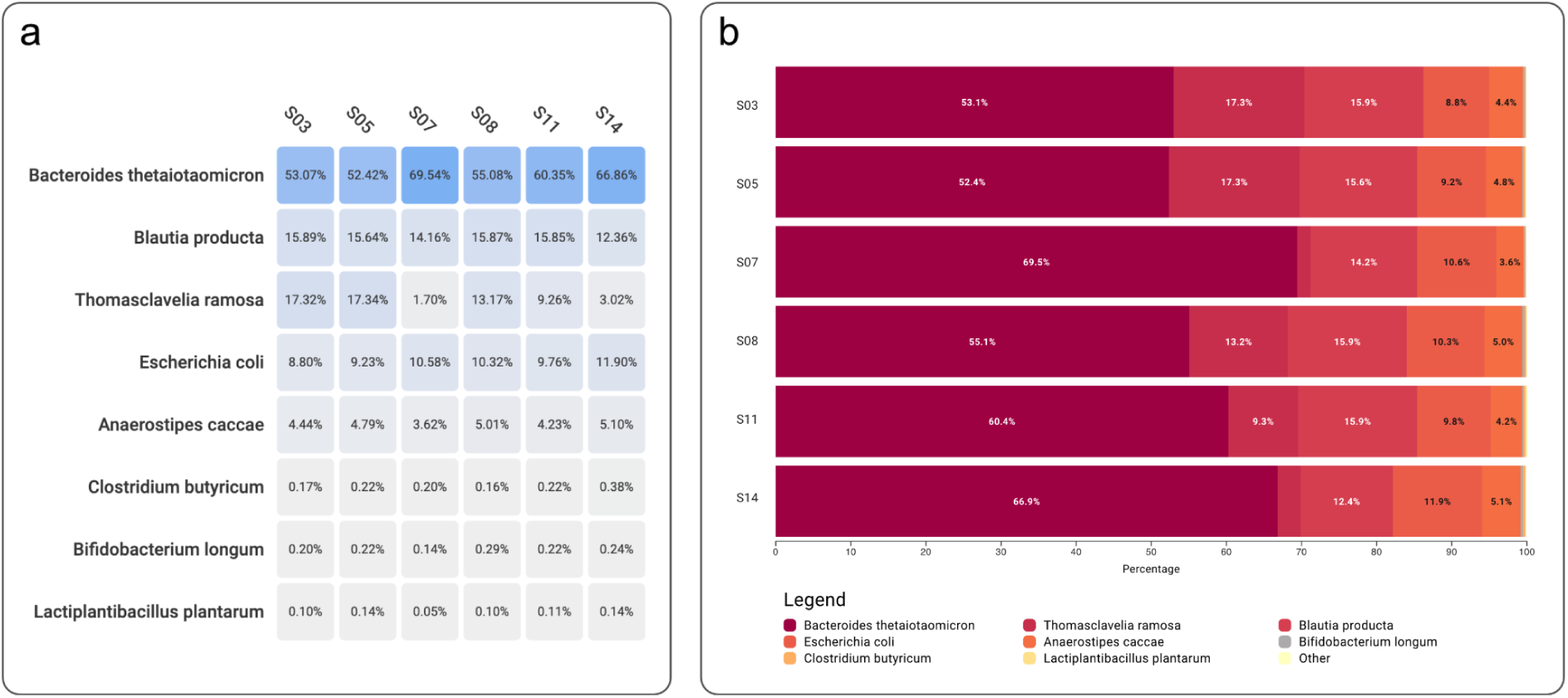
Taxonomic comparison of the six SIHUMIx samples in the Unipept web application, using the eight-species custom database. (a) Interactive heatmap of species-level assignments. Each cell gives the number of peptides whose lowest common ancestor is that species, as a percentage of all peptides assigned to one of the eight species in that sample, so each column sums to 100%. Peptides assigned at higher taxonomic ranks are not counted. (b) Comparative barplot of the same species-level assignments, showing the composition of each sample as stacked proportions.

*T. ramosa* varied most, from 1.7% in S07 to 17.3% in S03 and S05. Its three highest shares (S03, S05, S08) were all in samples that CAMPI identified as processed with more complex workflows. Because all six datasets derive from the same SIHUMIx culture, this variation most likely reflects differences between laboratory workflows rather than between communities.

For functional comparison, we launched PathwayPilot from the web interface, which placed the EC numbers of all six samples onto KEGG metabolic pathways, so each function appears in its metabolic context instead of in a flat list (Figure 2).

**Figure 2:**
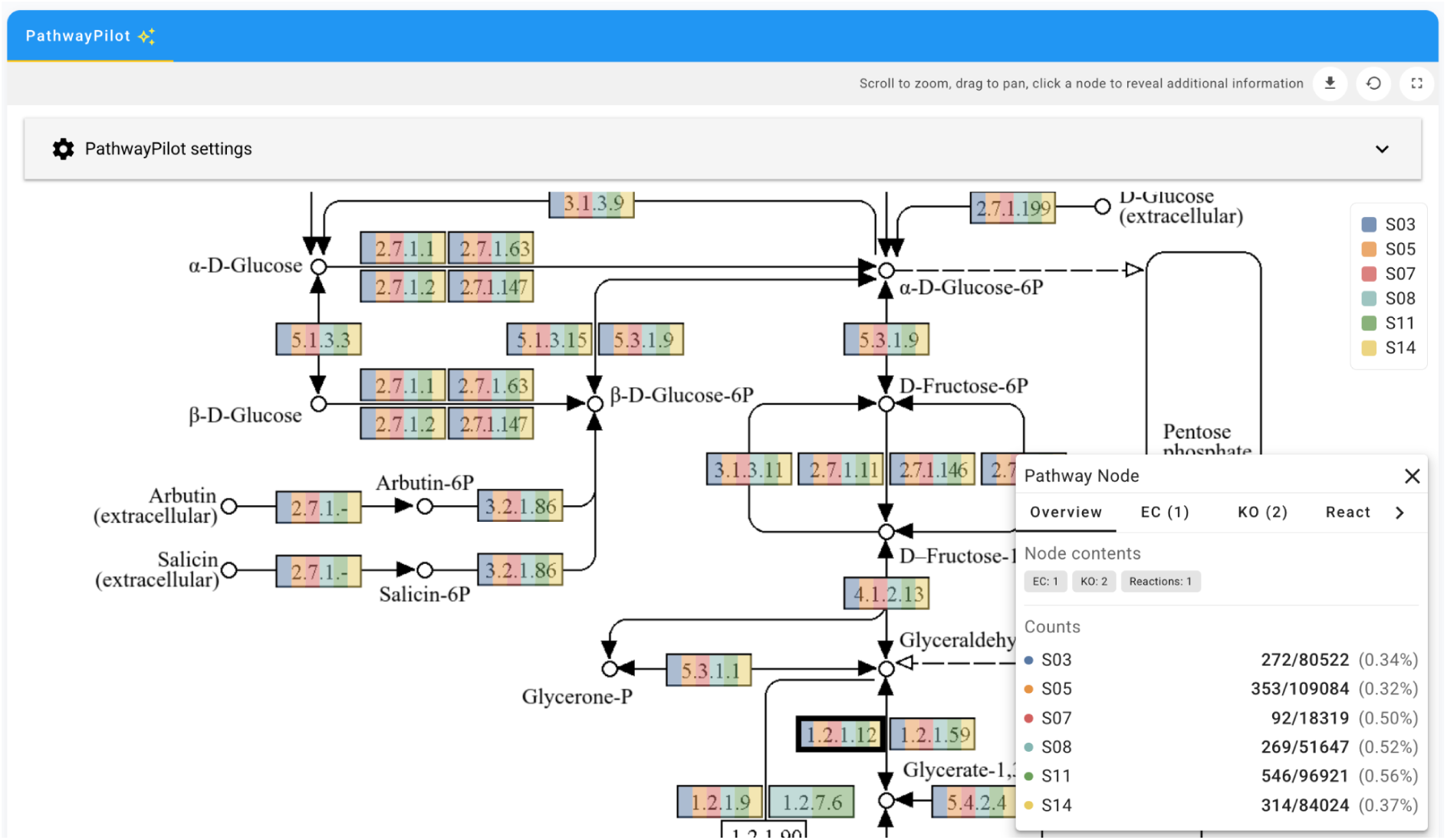
PathwayPilot view of KEGG glycolysis/gluconeogenesis (map00010) for the six SIHUMIx samples (S03, S05, S07, S08, S11, and S14). Each enzyme node is colored by the samples in which it was detected. The panel shows the peptide-spectrum matches per sample for the selected glyceraldehyde-3-phosphate dehydrogenase node (EC 1.2.1.12).

In glycolysis, each of the nine enzymatic steps along the central route from glucose 6-phosphate to pyruvate was detected in all six samples, with at least 32 peptide-spectrum matches per step and sample (**Table S2**). The pathway is therefore covered consistently across the different laboratory workflows, in line with CAMPI, which found the functional profiles of these samples highly similar based on their GO terms. In addition, each enzyme node shows its counts per sample and the taxa of its supporting peptides, as illustrated for glyceraldehyde-3-phosphate dehydrogenase (EC 1.2.1.12) in **Figure 2**. Such per-enzyme detail is not available from an overall similarity score.

### Backend throughput and deployability

We benchmarked query throughput for the three loading modes against the UniProtKB 2026_02 index, from unconstrained memory down to a 60 GB ceiling. All benchmarks ran on one virtual machine with 12 virtual CPUs (Intel Xeon Silver 4410Y host), 295 GB of RAM, and Ubuntu 24.04 (Linux kernel 6.8); the index was stored on a SATA SSD-backed virtual disk. Memory ceilings were enforced with Linux control groups (cgroups). Each point in **Figure 3** is the median throughput over 100 runs of 10 000 mixed-peptide queries, with peptide lengths drawn uniformly between 5 and 50 amino acids. All peptides were cut from UniProtKB proteins, so every query matched at least one protein. Under a memory ceiling, the server raised the number of worker threads from the default of 12, one per virtual CPU, to 96, to overlap page-fault latency with query work.

**Figure 3:**
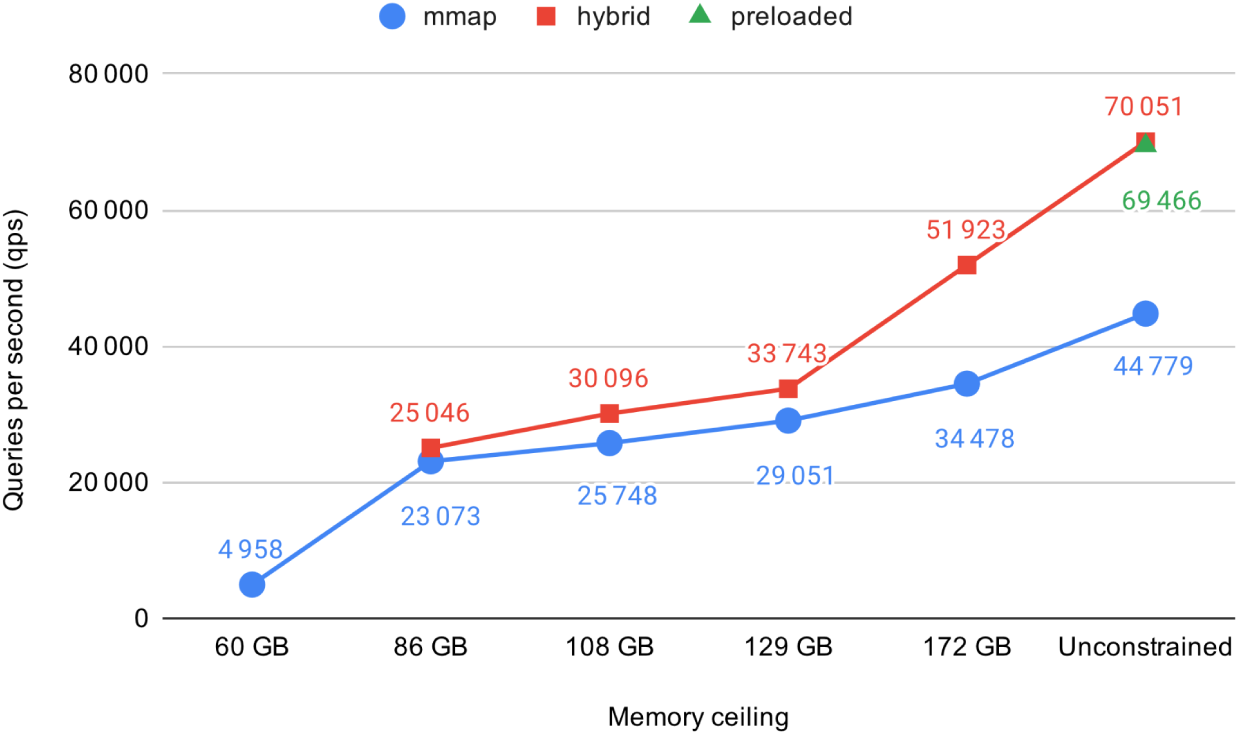
Median query throughput of the three index-loading modes as available RAM falls. The hybrid mode leads at every ceiling where it fits and still runs at 86 GB, where the fully preloaded index does not fit. At 60 GB, only memory-mapped loading runs. Under a memory ceiling, the server used 96 worker threads instead of the default 12.

Without a memory ceiling, the optimized index sustained about 69 500 queries per second in preloaded mode and 70 100 in the production hybrid mode (**Table 2**), equal within noise and both about 1.9 times the pre-optimization baseline of 37 500 queries per second. Memory-mapped loading reached 44 800 queries per second, about two-thirds of the hybrid throughput but still above the baseline. Cold starts were far shorter as well, at 3.2 s under mmap and 126 s under hybrid, against roughly 1 790 s (about 30 minutes) for the baseline.

**Table 2:** Deployment characteristics of the pre-optimization baseline and the three index-loading modes, from the same benchmark on the UniProtKB 2026_02 index (172 GB on disk). Throughput is the median number of queries per second (qps) at default threads without a memory ceiling; cold start is the time from server start to the first answered query.

| **Mode** | **Throughput** | **Cold start** | **Runs under a memory ceiling** | **When to use** |
|---|---|---|---|---|
| Baseline | 37 501 qps | 1 787 s | No | - |
| Preloaded | 69 466 qps | 392 s | No | Reference only; hybrid matches it with less memory |
| Hybrid | 70 051 qps | 126 s | Yes | Preloaded throughput at lower memory |
| mmap | 44 779 qps | 3.2 s | Yes | Memory is tight, or fast restart matters |

As memory tightens, throughput falls for every mode, and the fully preloaded index, which occupies roughly 186 GB in memory, stops fitting once the ceiling drops. Among the modes that still run, the hybrid mode leads at every ceiling where it fits and stays usable down to 86 GB, serving 25 046 queries per second against 23 073 for mmap (Figure 3). At 60 GB, only memory-mapped loading still ran, at 4 958 queries per second. The hybrid mode lowers the memory requirement for deployment, with a corresponding reduction in throughput under tighter memory limits.

## Conclusion

Since the 2024 update, Unipept has expanded both its analytical capabilities and the ways researchers can access them. The web application now combines project management, targeted reference selection, comparative visualization, and probabilistic inference, while project export supports sharing and later inspection of analyses. The Peptonizer2000, accelerated by the NORI inference engine, attaches a confidence score to each taxonomic and functional result, and an optional contaminant filter removes common laboratory contaminants before analysis. The SIHUMIx example illustrates how these capabilities can be used together in a single workflow.

Backend optimizations roughly doubled query throughput, and the hybrid loading mode provides deployment options for different memory budgets. The rewritten CLI extends programmatic access and reports the UniProtKB release used for analysis. Together, these developments support both interactive exploration and automated metaproteomics workflows.

Looking ahead, we plan to build on the project structure introduced here. Statistical comparison between groups of samples, such as treatment and control, would show which taxa and functions differ significantly instead of leaving this to visual inspection. Because such tests benefit from abundance information, Unipept will also accept quantitative input, such as spectral counts or intensities per peptide. Now that NORI makes the Peptonizer2000 fast enough for routine use, its confidence scores can also be shown in the comparative views, where they help weigh differences between samples by the strength of their peptide evidence. Finally, applying these analyses to studies with genuine biological variation, rather than a controlled community such as SIHUMIx, will test their value in practice.

## Data availability statement

Unipept is open-source software, and its source code is available in the Unipept GitHub organization (https://github.com/unipept): the web application (unipept), API server (unipept-api), SSA index (unipept-index), database construction scripts (unipept-database), command-line interface (unipept-cli), NORI inference engine (NORI), Peptonizer2000 (Peptonizer2000), and PathwayPilot (pathway-pilot). All repositories are released under the MIT license, except the Peptonizer2000 and PathwayPilot, which use the Apache 2.0 license. All analyses were performed with version 6.5.4 of the Unipept web application (unipept.ugent.be) against UniProtKB release 2026_02. The SIHUMIx datasets S03, S05, S07, S08, S11, and S14 are available from PRIDE under accession PXD023217, from which the X!Tandem results against the UniProt reference proteome database (files ending in_DB1UNIPROT.xml) were used. The filtered peptide lists, the script used to generate them, and the complete Unipept project are provided as Supporting Information (**Files S1 and S2**).

## Acknowledgements

This work was funded by the Research Foundation Flanders (FWO) [TVDB, LM: 1286824N and G010023N], Research Foundation Flanders (FWO) for ELIXIR Belgium [SVDV: I002819N], and Ghent University [PV: BOF/01P10623]. This project has received funding from the European Union’s Marie Skłodowska-Curie Actions (MSCA) Doctoral Networks under Grant Agreement No. 101225682 — METAMIC3.

## References

(1) Van Den Bossche, T.; Armengaud, J.; Benndorf, D.; Blakeley-Ruiz, J. A.; Brauer, M.; Cheng, K.; Creskey, M.; Figeys, D.; Grenga, L.; Griffin, T. J.; Henry, C.; Hettich, R. L.; Holstein, T.; Jagtap, P. D.; Jehmlich, N.; Kim, J.; Kleiner, M.; Kunath, B. J.; Malliet, X.; Martens, L.; Mehta, S.; Mesuere, B.; Ning, Z.; Tanca, A.; Uzzau, S.; Verschaffelt, P.; Wang, J.; Wilmes, P.; Zhang, X.; Zhang, X.; Li, L.; Metaproteomics Initiative. The Microbiologist’s Guide to Metaproteomics. iMeta 2025, 4 (3), e70031. 10.1002/imt2.70031

(2) Kleiner, M. Metaproteomics: Much More than Measuring Gene Expression in Microbial Communities. mSystems 2019, 4. 10.1128/mSystems.00115-19

(3) Salvato, F.; Hettich, R. L.; Kleiner, M. Five Key Aspects of Metaproteomics as a Tool to Understand Functional Interactions in Host-Associated Microbiomes. PLoS Pathog. 2021, 17 (2), e1009245. 10.1371/journal.ppat.1009245

(4) Sun, Z.; Ning, Z.; Figeys, D. The Landscape and Perspectives of the Human Gut Metaproteomics. Mol. Cell. Proteomics MCP 2024, 23 (5), 100763. 10.1016/j.mcpro.2024.100763

(5) Mesuere, B.; Devreese, B.; Debyser, G.; Aerts, M.; Vandamme, P.; Dawyndt, P. Unipept: Tryptic Peptide-Based Biodiversity Analysis of Metaproteome Samples. J. Proteome Res. 2012, 11 (12), 5773–5780. 10.1021/pr300576s

(6) Verschaffelt, P.; Tanca, A.; Abbondio, M.; Van Den Bossche, T.; Moortele, T. V.; Dawyndt, P.; Martens, L.; Mesuere, B. Unipept Desktop 2.0: Construction of Targeted Reference Protein Databases for Metaproteogenomics Analyses. J. Proteome Res. 2023, 22 (8), 2620–2628. 10.1021/acs.jproteome.3c00091

(7) Verschaffelt, P.; Van Thienen, P.; Van Den Bossche, T.; Van der Jeugt, F.; De Tender, C.; Martens, L.; Dawyndt, P.; Mesuere, B. Unipept CLI 2.0: Adding Support for Visualizations and Functional Annotations. Bioinformatics 2020, 36 (14), 4220–4221. 10.1093/bioinformatics/btaa553

(8) Mesuere, B.; Debyser, G.; Aerts, M.; Devreese, B.; Vandamme, P.; Dawyndt, P. The Unipept Metaproteomics Analysis Pipeline. PROTEOMICS 2015, 15 (8), 1437–1442. 10.1002/pmic.201400361

(9) The UniProt Consortium; Bateman, A.; Martin, M.-J.; Orchard, S.; Magrane, M.; Adesina, A.; Ahmad, S.; Bowler-Barnett, E. H.; Bye-A-Jee, H.; Carpentier, D.; Denny, P.; Fan, J.; Garmiri, P.; Gonzales, L. J. D. C.; Hussein, A.; Ignatchenko, A.; Insana, G.; Ishtiaq, R.; Joshi, V.; Jyothi, D.; Kandasaamy, S.; Lock, A.; Luciani, A.; Luo, J.; Lussi, Y.; Marin, J. S. M.; Raposo, P.; Rice, D. L.; Santos, R.; Speretta, E.; Stephenson, J.; Totoo, P.; Tyagi, N.; Urakova, N.; Vasudev, P.; Warner, K.; Wijerathne, S.; Yu, C. W.-H.; Zaru, R.; Bridge, A. J.; Aimo, L.; Argoud-Puy, G.; Auchincloss, A. H.; Axelsen, K. B.; Bansal, P.; Baratin, D.; Batista Neto, T. M.; Blatter, M.-C.; Bolleman, J. T.; Boutet, E.; Breuza, L.; Gil, B. C.; Casals-Casas, C.; Echioukh, K. C.; Coudert, E.; Cuche, B.; De Castro, E.; Estreicher, A.; Famiglietti, M. L.; Feuermann, M.; Gasteiger, E.; Gaudet, P.; Gehant, S.; Gerritsen, V.; Gos, A.; Gruaz, N.; Hulo, C.; Hyka-Nouspikel, N.; Jungo, F.; Kerhornou, A.; Mercier, P. L.; Lieberherr, D.; Masson, P.; Morgat, A.; Paesano, S.; Pedruzzi, I.; Pilbout, S.; Pourcel, L.; Poux, S.; Pozzato, M.; Pruess, M.; Redaschi, N.; Rivoire, C.; Sigrist, C. J. A.; Sonesson, K.; Sundaram, S.; Sveshnikova, A.; Wu, C. H.; Arighi, C. N.; Chen, C.; Chen, Y.; Huang, H.; Laiho, K.; Lehvaslaiho, M.; McGarvey, P.; Natale, D. A.; Ross, K.; Vinayaka, C. R.; Wang, Y.; Zhang, J. UniProt: The Universal Protein Knowledgebase in 2025. Nucleic Acids Res. 2025, 53 (D1), D609–D617. 10.1093/nar/gkae1010

(10) Vande Moortele, T.; Devlaminck, B.; Van de Vyver, S.; Van Den Bossche, T.; Martens, L.; Dawyndt, P.; Mesuere, B.; Verschaffelt, P. Unipept in 2024: Expanding Metaproteomics Analysis with Support for Missed Cleavages and Semitryptic and Nontryptic Peptides. J. Proteome Res. 2025, 24 (2), 949–954. 10.1021/acs.jproteome.4c00848

(11) Holstein, T.; Verschaffelt, P.; Van Den Bossche, T.; Van de Vyver, S.; Martens, L.; Mesuere, B.; Muth, T. The Peptonizer2000: Bringing Confidence to Metaproteomics. J. Proteome Res. 2026, 25 (4), 1831–1845. 10.1021/acs.jproteome.5c00567

(12) Van de Vyver, S.; Vande Moortele, T.; Binke, B.-B.; Verschaffelt, P.; Dawyndt, P.; Mesuere, B. NORI: Fast Probabilistic Inference for Ambiguous Observation-Entity Mappings. arXiv May 12, 2026. 10.48550/arXiv.2605.11648

(13) Vande Moortele, T.; Verschaffelt, P.; Huang, Q.; Doncheva, N. T.; Holstein, T.; Jachmann, C.; Dawyndt, P.; Martens, L.; Mesuere, B.; Van Den Bossche, T. PathwayPilot: A User-Friendly Tool for Visualizing and Navigating Metabolic Pathways. Mol. Cell. Proteomics 2025, 24 (3). 10.1016/j.mcpro.2025.100918

(14) Kanehisa, M.; Goto, S. KEGG: Kyoto Encyclopedia of Genes and Genomes. Nucleic Acids Res. 2000, 28 (1), 27–30. 10.1093/nar/28.1.27

(15) Van Den Bossche, T.; Kunath, B. J.; Schallert, K.; Schäpe, S. S.; Abraham, P. E.; Armengaud, J.; Arntzen, M. Ø.; Bassignani, A.; Benndorf, D.; Fuchs, S.; Giannone, R. J.; Griffin, T. J.; Hagen, L. H.; Halder, R.; Henry, C.; Hettich, R. L.; Heyer, R.; Jagtap, P.; Jehmlich, N.; Jensen, M.; Juste, C.; Kleiner, M.; Langella, O.; Lehmann, T.; Leith, E.; May, P.; Mesuere, B.; Miotello, G.; Peters, S. L.; Pible, O.; Queiros, P. T.; Reichl, U.; Renard, B. Y.; Schiebenhoefer, H.; Sczyrba, A.; Tanca, A.; Trappe, K.; Trezzi, J.-P.; Uzzau, S.; Verschaffelt, P.; von Bergen, M.; Wilmes, P.; Wolf, M.; Martens, L.; Muth, T. Critical Assessment of MetaProteome Investigation (CAMPI): A Multi-Laboratory Comparison of Established Workflows. Nat. Commun. 2021, 12 (1), 7305. 10.1038/s41467-021-27542-8

(16) Federhen, S. The NCBI Taxonomy Database. Nucleic Acids Res. 2012, 40 (Database issue), D136–143. 10.1093/nar/gkr1178

(17) Webb, E. Enzyme Nomenclature 1992. Recommendations of the Nomenclature Committee of the International Union of Biochemistry and Molecular Biology on the Nomenclature and Classification of Enzymes; Academic Press: San Diego, 1992.

(18) Ashburner, M.; Ball, C. A.; Blake, J. A.; Botstein, D.; Butler, H.; Cherry, J. M.; Davis, A. P.; Dolinski, K.; Dwight, S. S.; Eppig, J. T.; Harris, M. A.; Hill, D. P.; Issel-Tarver, L.; Kasarskis, A.; Lewis, S.; Matese, J. C.; Richardson, J. E.; Ringwald, M.; Rubin, G. M.; Sherlock, G. Gene Ontology: Tool for the Unification of Biology. Nat. Genet. 2000, 25 (1), 25–29. 10.1038/75556

(19) The Gene Ontology Consortium; Aleksander, S. A.; Balhoff, J.; Carbon, S.; Cherry, J. M.; Drabkin, H. J.; Ebert, D.; Feuermann, M.; Gaudet, P.; Harris, N. L.; Hill, D. P.; Lee, R.; Mi, H.; Moxon, S.; Mungall, C. J.; Muruganugan, A.; Mushayahama, T.; Sternberg, P. W.; Thomas, P. D.; Van Auken, K.; Ramsey, J.; Siegele, D. A.; Chisholm, R. L.; Fey, P.; Aspromonte, M. C.; Nugnes, M. V.; Quaglia, F.; Tosatto, S.; Giglio, M.; Nadendla, S.; Antonazzo, G.; Attrill, H.; dos Santos, G.; Marygold, S.; Strelets, V.; Tabone, C. J.; Thurmond, J.; Zhou, P.; Ahmed, S. H.; Asanitthong, P.; Luna Buitrago, D.; Erdol, M. N.; Gage, M. C.; Ali Kadhum, M.; Li, K. Y. C.; Long, M.; Michalak, A.; Pesala, A.; Pritazahra, A.; Saverimuttu, S. C. C.; Su, R.; Thurlow, K. E.; Lovering, R. C.; Logie, C.; Oliferenko, S.; Blake, J.; Christie, K.; Corbani, L.; Dolan, M. E.; Drabkin, H. J.; Hill, D. P.; Ni, L.; Sitnikov, D.; Smith, C.; Cuzick, A.; Seager, J.; Cooper, L.; Elser, J.; Jaiswal, P.; Gupta, P.; Jaiswal, P.; Naithani, S.; Lera-Ramirez, M.; Rutherford, K.; Wood, V.; De Pons, J. L.; Dwinell, M. R.; Hayman, G. T.; Kaldunski, M. L.; Kwitek, A. E.; Laulederkind, S. J. F.; Tutaj, M. A.; Vedi, M.; Wang, S.-J.; D’Eustachio, P.; Aimo, L.; Axelsen, K.; Bridge, A.; Hyka-Nouspikel, N.; Morgat, A.; Aleksander, S. A.; Cherry, J. M.; Engel, S. R.; Karra, K.; Miyasato, S. R.; Nash, R. S.; Skrzypek, M. S.; Weng, S.; Wong, E. D.; Bakker, E.; Berardini, T. Z.; Reiser, L.; Auchincloss, A.; Axelsen, K.; Argoud-Puy, G.; Blatter, M.-C.; Boutet, E.; Breuza, L.; Bridge, A.; Casals-Casas, C.; Coudert, E.; Estreicher, A.; Livia Famiglietti, M.; Feuermann, M.; Gos, A.; Gruaz-Gumowski, N.; Hulo, C.; Hyka-Nouspikel, N.; Jungo, F.; Le Mercier, P.; Lieberherr, D.; Masson, P.; Morgat, A.; Pedruzzi, I.; Pourcel, L.; Poux, S.; Rivoire, C.; Sundaram, S.; Bateman, A.; Bowler-Barnett, E.; Bye-A-Jee, H.; Denny, P.; Ignatchenko, A.; Ishtiaq, R.; Lock, A.; Lussi, Y.; Magrane, M.; Martin, M. J.; Orchard, S.; Raposo, P.; Speretta, E.; Tyagi, N.; Warner, K.; Zaru, R.; Diehl, A. D.; Lee, R.; Chan, J.; Diamantakis, S.; Raciti, D.; Zarowiecki, M.; Fisher, M.; James-Zorn, C.; Ponferrada, V.; Zorn, A.; Ramachandran, S.; Ruzicka, L.; Westerfield, M. The Gene Ontology Knowledgebase in 2023. Genetics 2023, 224 (1), iyad031. 10.1093/genetics/iyad031

(20) Paysan-Lafosse, T.; Blum, M.; Chuguransky, S.; Grego, T.; Pinto, B. L.; Salazar, G. A.; Bileschi, M. L.; Bork, P.; Bridge, A.; Colwell, L.; Gough, J.; Haft, D. H.; Letunić, I.; Marchler-Bauer, A.; Mi,H.; Natale, D. A.; Orengo, C. A.; Pandurangan, A. P.; Rivoire, C.; Sigrist, C. J. A.; Sillitoe, I.; Thanki, N.; Thomas, P. D.; Tosatto, S. C. E.; Wu, C. H.; Bateman, A. InterPro in 2022. Nucleic Acids Res. 2023, 51 (D1), D418–D427. 10.1093/nar/gkac993

(21) Sutton, C.; McCallum, A. Improved Dynamic Schedules for Belief Propagation. In Proceedings of the Twenty-Third Conference on Uncertainty in Artificial Intelligence; UAI’07; AUAI Press: Arlington, Virginia, USA, 2007; pp 376–383.

(22) Knoll, C.; Rath, M.; Tschiatschek, S.; Pernkopf, F. Message Scheduling Methods for Belief Propagation. In Machine Learning and Knowledge Discovery in Databases; Appice, A., Rodrigues, P. P., Santos Costa, V., Gama, J., Jorge, A., Soares, C., Eds.; Springer International Publishing: Cham, 2015; pp 295–310. 10.1007/978-3-319-23525-7_18

(23) Krause, J. L.; Schaepe, S. S.; Fritz-Wallace, K.; Engelmann, B.; Rolle-Kampczyk, U.; Kleinsteuber, S.; Schattenberg, F.; Liu, Z.; Mueller, S.; Jehmlich, N.; Von Bergen, M.; Herberth, G. Following the Community Development of SIHUMIx – a New Intestinal in Vitro Model for Bioreactor Use. Gut Microbes 2020, 11 (4), 1116–1129. 10.1080/19490976.2019.1702431

(24) Perez-Riverol, Y.; Bandla, C.; Kundu, D. J.; Kamatchinathan, S.; Bai, J.; Hewapathirana, S.; John, N. S.; Prakash, A.; Walzer, M.; Wang, S.; Vizcaíno, J. A. The PRIDE Database at 20 Years: 2025 Update. Nucleic Acids Res. 2025, 53 (D1), D543–D553. 10.1093/nar/gkae1011

(25) Van Den Bossche, T.; Arntzen, M. Ø.; Becher, D.; Benndorf, D.; Eijsink, V. G. H.; Henry, C.; Jagtap, P. D.; Jehmlich, N.; Juste, C.; Kunath, B. J.; Mesuere, B.; Muth, T.; Pope, P. B.; Seifert, J.; Tanca, A.; Uzzau, S.; Wilmes, P.; Hettich, R. L.; Armengaud, J. The Metaproteomics Initiative: A Coordinated Approach for Propelling the Functional Characterization of Microbiomes. Microbiome 2021, 9 (1), 243. 10.1186/s40168-021-01176-w

(26) Van Den Bossche, T.; Grenga, L.; Alves, G.; Arntzen, M. Ø.; Benndorf, D.; Brauer, M.; Figeys, D.; Henry, C.; Hettich, R. L.; Heyer, R.; Jagtap, P. D.; Jehmlich, N.; Kleiner, M.; Li, L.; Mesuere, B.; Pabst, M.; Pandhal, J.; Pope, P. B.; Seifert, J.; Trautwein-Schult, A.; Verschaffelt, P.; Wilmes, P.; Armengaud, J.; Kunath, B. J. The Metaproteomics Initiative: Five Years of Community-Driven Progress. Microbiome 2026, 14 (1), 186. 10.1186/s40168-026-02463-0

(27) Craig, R.; Beavis, R. C. TANDEM: Matching Proteins with Tandem Mass Spectra. Bioinformatics 2004, 20 (9), 1466–1467. 10.1093/bioinformatics/bth092

(28) Maturana, J. L.; Cárdenas, J. P. Insights on the Evolutionary Genomics of the Blautia Genus: Potential New Species and Genetic Content Among Lineages. Front. Microbiol. 2021, 12. 10.3389/fmicb.2021.660920

