## Supporting information for "Unipept in 2026: A Unified Web Platform for Comparative Metaproteomics Analysis"

### Contents

Table S1: Peptide filtering and matching for the six SIHUMIx samples
Table S2: Peptide-spectrum matches supporting the nine core steps of glycolysis
File S1: Unipept project file of the six SIHUMIx samples (separate file)
File S2: Filtered peptide lists and filtering script (separate file)

### Table S1

| Sample | Spectra (decoy) | Target PSMs retained | FDR | Unique peptides | Matched to UniProtKB | Matched to 8 species |
|---|---|---|---|---|---|---|
| S03 | 127 549 (13 625) | 80 522 | 0.32% | 27 187 | 27 187 (100.00%) | 27 066 (99.55%) |
| S05 | 194 141 (25 317) | 109 084 | 0.40% | 30 815 | 30 815 (100.00%) | 30 644 (99.45%) |
| S07 | 35 901 (6 614) | 18 319 | 0.58% | 13 391 | 13 391 (100.00%) | 13 307 (99.37%) |
| S08 | 73 110 (5 743) | 51 647 | 0.32% | 20 389 | 20 389 (100.00%) | 20 240 (99.27%) |
| S11 | 149 782 (18 182) | 96 921 | 0.54% | 35 349 | 35 348 (>99.99%) | 35 101 (99.30%) |
| S14 | 163 333 (25 043) | 84 024 | 0.43% | 19 297 | 19 295 (99.99%) | 19 146 (99.22%) |
| Total | 743 816 (94 524) | 440 517 | – | 146 428 | 146 425 (>99.99%) | 145 504 (99.37%) |

**Table S1:** Peptide filtering and matching for the six SIHUMIx samples. Counts for S11 and S14 were obtained after pooling their four fractions and three replicates, respectively. Spectra are those with a peptide assignment in the X!Tandem results, with the number whose best-scoring hit was a decoy in parentheses. Target peptide-spectrum matches (PSMs) retained are the target matches passing the CAMPI e-value threshold of 0.186, after removing spectra whose best-scoring hits were tied between different peptides; the false discovery rate (FDR) is the target–decoy FDR at that threshold. Unique peptides are the peptides retained after filtering. Percentages in the last two columns are relative to the unique peptides of each sample.

### Table S2

| Step | EC number(s) | S03 | S05 | S07 | S08 | S11 | S14 |
|---|---|---|---|---|---|---|---|
| Glucose-6-phosphate isomerase | 5.3.1.9 | 202 | 255 | 65 | 128 | 318 | 241 |
| Phosphofructokinase | 2.7.1.11, 2.7.1.90 | 450 | 572 | 89 | 239 | 392 | 549 |
| Fructose-bisphosphate aldolase | 4.1.2.13 | 1 122 | 1 505 | 145 | 391 | 781 | 842 |
| Triosephosphate isomerase | 5.3.1.1 | 398 | 602 | 58 | 224 | 251 | 404 |
| Glyceraldehyde-3-phosphate dehydrogenase | 1.2.1.12, 1.2.1.59 | 248 | 327 | 83 | 197 | 478 | 294 |
| Phosphoglycerate kinase | 2.7.2.3 | 776 | 1 047 | 138 | 376 | 635 | 675 |
| Phosphoglycerate mutase | 5.4.2.11, 5.4.2.12 | 200 | 242 | 38 | 153 | 146 | 247 |
| Enolase | 4.2.1.11 | 281 | 373 | 65 | 184 | 276 | 687 |
| Pyruvate kinase | 2.7.1.40 | 203 | 284 | 32 | 143 | 201 | 162 |

**Table S2:** Peptide-spectrum matches supporting the nine core steps of glycolysis (KEGG map00010) in the six SIHUMIx samples. A peptide was counted for a step if Unipept annotated it with one of the listed Enzyme Commission (EC) numbers and more than half of its matched UniProtKB proteins carry the name of that enzyme. Steps with two EC numbers are catalyzed by alternative enzymes, which are counted together.

### Supporting files

**File S1** (.zip). Unipept project containing the six SIHUMIx samples, their analysis settings, the eight-species custom database, and the results. The .unipept file inside can be imported into the Unipept web application (https://unipept.ugent.be) to inspect all analyses described in the main text.

**File S2** (.zip). Filtered peptide lists of the six SIHUMIx samples (one peptide per peptide-spectrum match), and the script used to derive them from the X!Tandem results in PRIDE (PXD023217).
